# projectR: An R/Bioconductor package for transfer learning via PCA, NMF, correlation, and clustering

**DOI:** 10.1101/726547

**Authors:** Gaurav Sharma, Carlo Colantuoni, Loyal A Goff, Elana J Fertig, Genevieve Stein-O’Brien

## Abstract

**Motivation:** Dimension reduction techniques are widely used to interpret high-dimensional biological data. Features learned from these methods are used to discover both technical artifacts and novel biological phenomena. Such feature discovery is critically import to large single-cell datasets, where lack of a ground truth limits validation and interpretation. Transfer learning (TL) can be used to relate the features learned from one source dataset to a new target dataset to perform biologically-driven validation by evaluating their use in or association with additional sample annotations in that independent target dataset.

**Results:** We developed an R/Bioconductor package, projectR, to perform TL for analyses of genomics data via TL of clustering, correlation, and factorization methods. We then demonstrate the utility TL for integrated data analysis with an example for spatial single-cell analysis.

**Availability:** projectR is available on Bioconductor and at https://github.com/genesofeve/projectR.

## Introduction

Dimension reduction methods play a key role in biological discovery from high-dimensional genomics datasets. The lower-dimensional spaces learned represent both biological information and technical artifacts. Thus, it is crucial to interpret and validate these spaces. Independent datasets from related but varied biological contexts, such as different data modalities of equivalent samples or data from the same tissue in related organisms, can be used for interpretation and validation as only the biological effects, and not the technical effects, will be shared. Thus, we can use transfer learning (TL), a sub-domain of machine learning, for *in silico* validation, interpretation, and exploration of these spaces using independent but related datasets (Stein-O’Brien *et al.*, 2019; Taroni *et al.*, 2019). Furthermore, once the robustness of biological signal is established, these TL approaches can be used for multimodal data integration (Stuart *et al.*, 2019). Here, we develop the projectR package to perform TL for dimension reduction techniques for genomics analysis.

## Methods

The projectR package performs TL from the outputs of PCA (Principal Component Analysis), NMF (Non-negative Matrix Factorization), regression, K-means, hierarchical clustering, and correlation via the main function of the package--projectR. The inputs to projectR are target data--data--and learned gene features--loadings. To match genes between the datasets, the software contains a function, geneMatchR, which returns the two input datasets separately or jointly, if merge is true, with only common genes. Utilizing an S4 generic, projectR’s loadings argument corresponds to features for classes prcomp, LinearEmbeddingMatrix, matrix, kmeans, hclust, and correlateR (defined in the projectR package) for spaces learned by PCA, NMF, regression, K-means and hierarchical clustering, and correlation respectively (Meng *et al.*, 2016). projectR returns a matrix with sample weights for each input basis in the loadings matrix with the option to include p-values (Wald-test) for each value in the projection matrix (Supplemental File 1).

To facilitate TL further, additional functions are provided to operate on the output of projectR. Example uses include identifying the patterns that are predictive of sample annotations such as cell type or transferring annotations using previous associated patterns. The aucMat function identifies the patterns predictive of given sample annotations using the performance and prediction function from the ROCR package. The alluvialMat function generates an alluvial plot given projection matrix from projectR and annotations as input.

To demonstrate the application of projectR in spatial single-cell analysis we selected high-quality expression data of 1297 cells and 8097 genes from development stage 6 *Drosophila* embryo as source data generated by (Karaiskos *et al.*, 2017) available at https://shiny.mdc-berlin.de/DVEX/. The position of almost all of the fly embryo cells can be specified using the binarized expression from *in situ* imaging of 84 marker genes identified by Berkeley *Drosophila* Transcription Network Project (Karaiskos *et al.*, 2017; Fowlkes *et al.*, 2008). The code for this analysis is available at https://github.com/fertigLab/projectRSpatialExample. We validated the patterns by comparing them to spatial patterns given with vISH (virtual *in-situ* hybridization) (https://shiny.mdc-berlin.de/DVEX/) computed by DistMap (Karaiskos *et al.*, 2017).

## Results

ProjectR perform TL on gene signatures from clustering, PCA, NMF, and correlation. It is computationally fast taking 8.09 ± 0.51 s on a 16 GB, Intel Core i7-8750H based 64-bit Windows 10 computer for projecting a 20000×1000 target dataset on 20000×100 latent space. Previously, we demonstrated the ability of this approach to relate molecular signatures associated with retinal development with our Bayesian NMF algorithm CoGAPS (Fertig et al., 2010) across data platforms, tissues, and species (Stein-O’Brien et al., 2019). To further demonstrate the utility of this approach we apply projectR for multi-modal data integration to enable spatial single-cell analysis. After analysis with CoGAPS (Fertig et al., 2010) on scRNAseq from Drosophila embryo, we input these patterns as the loadings and the binarized gene-marker by position matrix from the in situ imaging as the data (see methods). Figure 1 demonstrates that this analysis can transfer the full molecular states from the non-spatially resolved single-cell data onto the spatially resolved imaging data. In addition to their spatial resolution, we find that the top five genes that distinguish developmental gradients from the TL are also drivers of developmental gradients in the Drosophila. For example, we found the top genes associated with the ventral pattern (Fig 1(b)) were: Ilp4, twi, Cyp310a1, ventrally-expressed-protein-D, and CG4500 and all of them were predominantly ventrally expressed as confirmed by vISH (Karaiskos et al., 2017; Fowlkes et al., 2008).

**Fig 1.**
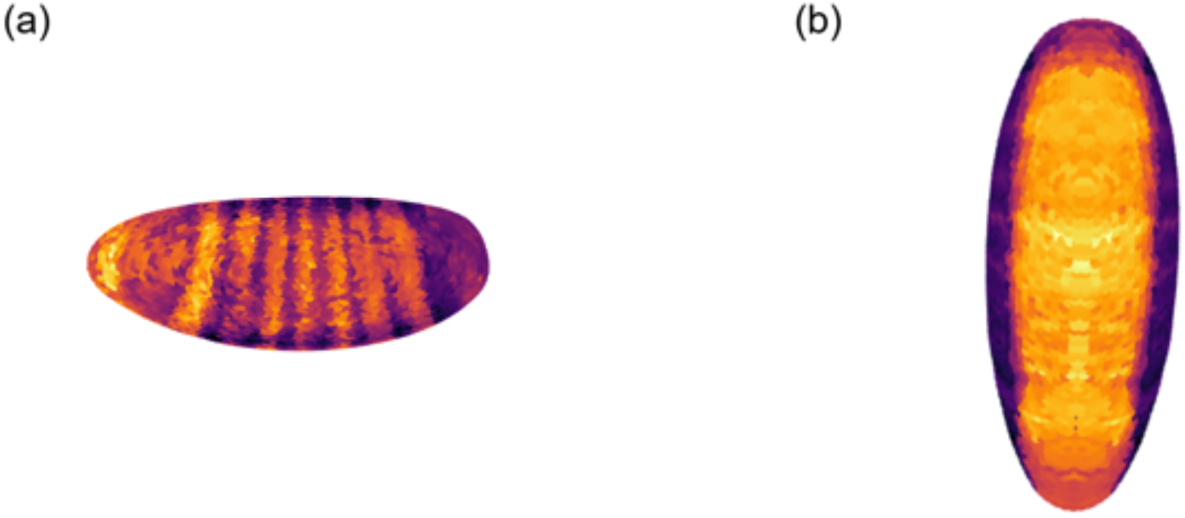
Spatial gene expression patterns identified in *Drosophila* stage 6 embryo using projectR and CoGAPS. (a) Anterior-posterior gene expression pattern characteristic of fly development. (b) A pattern shown by predominantly ventrally expressed genes.

## Discussion

We developed projectR as a software package to enable TL dimension reduction of genomics data. We previously showed that application of this technique to patterns learned from NMF relates datasets from different species, data modalities, tissues, and measurement platforms (Stein-O’Brien *et al.*, 2019). In this paper, we demonstrate its further utility to integrate imaging and single-cell data for spatial transcriptional analysis and expansion to dimension reduction techniques beyond NMF. While similar to Slide-seq, we note projectR generalizes beyond a NMF-based regression framework to implement high spatial resolution of transcriptional data (Rodriques *et al.*, 2019). The software is developed generally to enable pattern validation, discovery, and annotation transfer across datasets with a wide range of unsupervised learning techniques.

## Acknowledgements

We thank Timothy Triche, Jr and Thomas Sherman for feedback.

