## Supplemental File 1 for "projectR: An R/Bioconductor package for transfer learning via PCA, NMF, correlation, and clustering"

### Projection calculation

Here we describe projectR using gene expression data as an example. Given a target data matrix $T$ (genes by samples), loadings matrix $L$ (genes by patterns) learned from the source data, we can compute a projection matrix $P$ by projecting $T$ onto low-dimensional latent space defined by $L$. A projection can be defined as a mapping of the points from one space to another generally lower-dimensional space. The mapping can be mathematically written as a function $\varphi(x) = y : \Re^D \mapsto \Re^d \text{ s.t. } d \leq D \text{ for } x \in \Re^D, y \in \Re^d$. Using the common genes in the source and target data, we employ different methods to calculate projection depending on the loadings. For our default method, we use multiple linear regressions to calculate pattern weights in $P$ using `lmFit` function of the `limma` package in R. Mathematically,

$$T_j \sim L\beta_j$$

where $T_j$ is the $j^{th}$ column of $T$, and $\beta_j$ is the $j^{th}$ column of P. It generates a least square fit, and the projection matrix is generated by taking orthogonal projections in the column space of $L$.

The projection weights were calculated using linear regression. For each pattern weight $\beta_{ij}$, we can test the null hypothesis $H_0 = \beta_{0ij}$ by computing t-statistic, $t_{ij}$,

$$t_{ij} = \frac{\beta_{ij} - \beta_{0ij}}{se(\beta_{ij})}$$

and check its significance using a t-distribution with $n - p$ degrees of freedom where $n$ is the number of genes and $p$ is the number of patterns.

For loadings calculated using PCA which gives us principal directions of variation given by the eigenvectors as axes and origin as the centroid of the data. To project a given target data matrix, T, we apply the same approach that is used in PCA to find new weights in the PC space. The first step is to center expression for each gene around its mean, that is, subtract the mean gene expression from for all the genes. $g_{ij} = g_{ij} - \bar{g}_j$ This gives as a centered target data matrix, $\bar{T}$. Using the eigenvectors obtained from the PCA of the source data as loadings $L$, the target data $T$ can be projected onto it giving the projection $P$ as $P = \bar{T}'L$

Similar to k-means clustering we can use `cutree` function in R to specify the number of clusters for hierarchical clustering. We create the loadings matrix $L$ with the number of columns equal to the number of clusters. For each gene present in a cluster, the entry is specified as its correlation with the mean expression across samples. Given the target data matrix $T$, $L$ can be used to obtain the projection using the default method.
